# Purkinje Cell Activity in Medial and Lateral Cerebellum During Suppression of Voluntary Eye Movements in Rhesus Macaques

**DOI:** 10.1101/2021.03.26.437236

**Authors:** Eric Avila, Nico A. Flierman, Peter J. Holland, Pieter R. Roelfsema, Maarten A. Frens, Aleksandra Badura, Chris I. De Zeeuw

## Abstract

Volitional suppression of responses to distracting external stimuli enables us to achieve our goals. This volitional inhibition of a specific behavior is supposed to be mainly mediated by the cerebral cortex. However, recent evidence supports the involvement of the cerebellum in this process. It is currently not known whether different parts of the cerebellar cortex play differential or synergistic roles in planning and execution of this behavior. Here, we measured Purkinje cell (PC) responses in the medial and lateral cerebellum in two rhesus macaques during a pro- and antisaccade task. During an antisaccade trial, non-human primates were instructed to make a saccadic eye movement away from a target, rather than towards it, as in prosaccade trials. Our data shows that the cerebellum plays an important role not only during execution of the saccades, but also during the volitional inhibition of eye movements towards the target. Simple Spike (SS) modulation during the instruction and execution period of pro- and antisaccades was prominent in PCs of both medial and lateral cerebellum. However, only the SS activity in the lateral cerebellar cortex contained information about trial identity and showed a stronger reciprocal interaction with complex spikes. Moreover, SS activity of different PC groups modulated bidirectionally in both regions, but the PCs that showed facilitating and suppressive activity were predominantly associated with instruction and execution, respectively. These findings show that different cerebellar regions and PC groups contribute to goal-directed behavior and volitional inhibition, but with different propensities, highlighting the rich repertoire of cerebellar control in executive functions.

**Significance Statement:** The antisaccade task is commonly used in research and clinical evaluation as a test of volitional and flexible control of behavior. It requires volitional suppression of prosaccades, a function that has been attributed to the neocortex. However, recent findings indicate that cerebellum also contributes to this behavior. We recorded from neurons in the medial and lateral cerebellum to evaluate their responses in this task. We found that both regions significantly modulated their activity during this task, but only cells in the lateral cerebellum encoded the stimulus identity in each trial. These results indicate that the cerebellum actively contributes to the control of flexible behavior and that lateral and medial cerebellum play different roles during volitional eye movements.

## Introduction

In a dynamic environment, volitional control of behavior is necessary to make flexible, well-adapted choices. This often requires selective suppression of responses to external stimuli, a hallmark of conscious, executive control (Diamond, 2013). In a laboratory setting, we can measure this complex behavior using the antisaccade task. It requires the subjects to refrain from looking at a suddenly appearing target and instead execute a saccade to the (unmarked) mirror position of that target (Everling and Fischer, 1998; Mitchell et al., 2002; Munoz and Everling, 2004). Similar to other volitional movements, many regions in the cerebral cortex have been identified as crucial for the correct execution of the antisaccade task (Funahashi et al., 1993; Schlag-Rey et al., 1997; Gottlieb and Goldberg, 1999; Bunge et al., 2005; Hakvoort Schwerdtfeger et al., 2012; Everling and Johnston, 2013; Cutsuridis et al., 2014). Given its reciprocal connections with the relevant neocortical regions (Kelly and Strick, 2003), the cerebellum may form an additional hub in this voluntary motor control circuitry (Tanaka et al., 2003; Peterburs et al., 2012; Brunamonti et al., 2014; Kunimatsu et al., 2016; Dacre et al., 2019). This possibility is corroborated by recent findings that the cerebellum participates in movement planning (Ashmore and Sommer, 2013; Giovannucci et al., 2017; Deverett et al., 2018; Gao et al., 2018; Kostadinov et al., 2019). However, to what extent different parts of the cerebellum contribute to the planning and execution of the antisaccades, and complex movements in general, is unclear (Miall et al., 1993; Thach, 2007; Ito, 2008; Gao et al., 2018, 2019; Chabrol et al., 2019; De Schutter, 2019).

Several studies have indicated that execution of simple or reflexive movements may be controlled by Purkinje cells (PCs) in the medial part of the cerebellum, whereas complex behaviors, which require instruction, inhibition and planning, might be controlled by more lateral parts (Strick et al., 2009; Caligiore et al., 2017; Chabrol et al., 2019; De Schutter, 2019; Sendhilnathan et al., 2020). Others however, advocate that both simple and complex movements can be controlled by the same group of PCs (Gao et al., 2018, 2019), the location of which in the cerebellar cortex may be determined by the PC-to-effector pathway (Voogd et al., 2012). Settling these questions would require monitoring of spiking activity of PCs in both medial and lateral cerebellar areas during a motor task that includes the execution of both simple and complex forms of movements related to the same effector.

Here, we investigated the hypotheses that (1) PCs in the cerebellum play a role in the volitional inhibition of stimulus-driven eye movements during the antisaccade task; and (2) that PCs in medial and lateral cerebellum differentially contribute to pro- and antisaccades. More specifically, we set out to study modulation of randomly selected saccade-related PCs in the medial (oculomotor vermis, OMV) and lateral (crus-I/II) cerebellum of non-human primates (macaca mulatta) during the generation of both pro- and antisaccades (**Fig. 1**). We found that Simple Spike (SS) modulation during the instruction and execution periods of pro- and antisaccade trials was prominent in PCs in both cerebellar regions. However, whereas SS activity in the lateral cerebellum contained information about the trial identity, we were not able to detect this in the vermis. In addition, Complex Spike (CS) activity showed a higher level of reciprocity with respect to SS activity in the lateral cerebellum. Although the PC activity modulated bidirectionally in both cerebellar regions, the PCs in the lateral cerebellum that facilitated their activity, so called upbound PCs (De Zeeuw, 2021), contributed most prominently to the instruction of antisaccade trials. Together, our data are adding to the growing body of evidence for a role of the cerebellum in executive control and are inciting the field to explore cerebellar activity under conditions where response inhibition is engaged.

**Figure 1.**
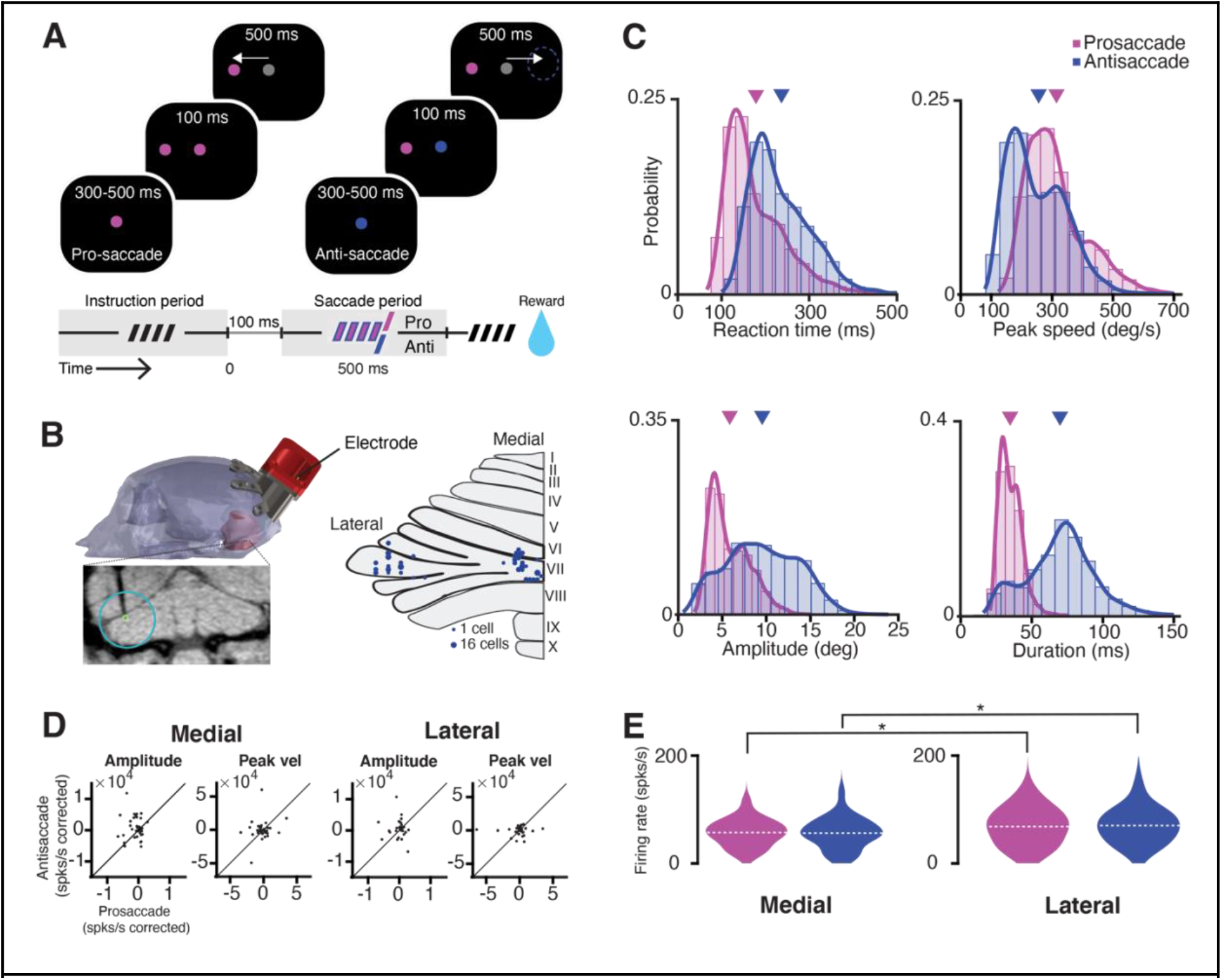
Task, recording location, saccade kinematics and related PC firing characteristics. ***A***, A scheme representing the different periods of the task. Trial condition was indicated by the color of the fixation point. After a random delay between 300-500 ms (i.e., instruction period, dashed symbol) a target appeared in one of 8 locations, while NHPs were required to maintain fixation for 100 ms. Next, the fixation target switched to gray, and NHPs were given 500 ms to execute a saccade (i.e., saccade period) towards the target (prosaccade) or the mirror position of the target (antisaccade). ***B, left***, CT-based 3D-reconstruction of the skull of Mi; ***bottom***, coronal MRI section showing the localization of an electrode in the left lateral cerebellum; ***right***, overview of the localization of recordings sites in the medial (vermal lobules VIc and VII) and lateral cerebellum (crus-I/II, left hemisphere). ***C***, Features of kinematics of all prosaccades (magenta; monkey Mo *n* = 6114, monkey Mi *n* = 6670) and antisaccades (blue; Mo *n* = 4114, Mi *n* = 6305) for all trials over the 122 recording sessions. Antisaccades were characterized by longer reaction times (pro-mean 178.4 ms ± 70.6 s.d.; anti-mean 236.6 ms ± 72.7 s.d.; *p* < 0.0001), lower peak velocities (pro-mean 311.8°/s ± 102.6 s.d.; anti-mean 253.4°/s ± 103.1 s.d.; *p* < 0.0001), bigger amplitudes (pro-mean 5.8° ± 2.3 s.d.; anti-mean 9.5° ± 4.1 s.d.; *p* < 0.0001), and longer durations (pro-mean 34.7 ms ± 7.9 s.d.; anti-mean 69.8 ms ± 24 s.d.; *p* < 0.0001). Triangles in magenta and purple indicate means of pro- and antisaccade movement parameters, respectively. ***D,*** Scatter plots with corrected PC activity after a linear regression model fitted to amplitude or peak speed during all pro- and antisaccades in medial (left) and lateral (right) cerebellum. ***E,*** Violin plots showing the mean average simple spike firing rate for all cells during prosaccades (magenta) and antisaccades (blue) for medial (left) and lateral (right) cerebellum). Dashed lines depict the mean value for the population. The asterisks show significance levels for a Wilcoxon rank sum test for the comparison between medial and lateral cerebellum in each condition (pro medial vs. lateral *p* = 0.04; anti medial vs. lateral *p* = 0.03).

## Materials and Methods

### Animals

All procedures complied with the NIH Guide for the Care and Use of Laboratory Animals (National Institutes of Health, Bethesda, Maryland), and were approved by the institutional animal care and use committee of the Royal Netherlands Academy of Arts and Sciences (AVD8010020184587). Two adult male non-human primates (NHPs, Macaca mulatta), Mi and Mo, were used in this study.

### Surgical procedures

Animals were prepared for eye movement and awake, extracellular single unit recordings in the cerebellum using surgical and electrophysiological techniques in a two-step procedure (Roelfsema et al., 2012). Under general anesthesia induced with ketamine (15 mg/kg, i.m.) and maintained under intubation by ventilating with a mixture of 70% N2O and 30% O2, supplemented with 0.8% isoflurane, fentanyl (0.005 mg/kg, i.v.), and midazolam (0.5 mg/kg • h, i.v.), we first implanted a titanium head holder to painlessly immobilize the NHP’s head. Four months later, once the NHPs had mastered the behavioral task (see below), a custom-made 40 mm chamber was implanted under the same anesthesia conditions as described above to gain access to the cerebellum with a 25 angle (**Fig. 1B**). Animals recovered for at least 21 days before training was resumed.

### Behavioral task

Animals were trained to perform a randomized interleaving pro- and antisaccade task in 8 different directions (cardinal and diagonal directions), with amplitudes between 5 ° and 14 ° (**Fig. 1A**). All recordings were conducted in complete darkness. During training and experiments, NHPs were seated in a primate chair (Crist Instrument, USA) with their head restrained at 100 cm from a screen with a resolution of 1,024 x 768 pixels. Visual stimuli were presented by a CRT-Projector Marquee 9500 LC (VDC Display Systems, Florida, USA) with a refresh rate of 100 Hz. Binocular vision was unrestricted. A trial started when the animal fixated on a red or green fixation point at the center of the screen for a random time between 300-500 ms. Next, a red target appeared in one of 8 different target locations. After 100 ms the fixation point changed to gray and the NHP was allowed to perform either a prosaccade toward the target (when fixation point was red) or antisaccade in the opposite direction with the same amplitude as the target, (when fixation point was green) within 500 ms (in the figures colors of fixation point and target were changed to blue and magenta to accommodate colorblind readers). The animals received liquid reward if they performed a correct saccade (within 6 ° of the (anti-)target) and maintained fixation for 100 ms. Within a single block the 8 (targets) * 2 (pro/anti) possible configurations were presented in a random order.

### Recordings

Position of the right eye was recorded with an infrared video-based eye tracker (iViewX Hi-Speed Primate, SMI GmbH, Germany) at a sampling rate of 350 Hz. The eye tracker was calibrated before every recording session by having the animal look at 1° target grid consisting of 9 points (one at the center of the screen and 8 points 10° apart) to adjust the offset of X and Y position by hand. Pre and post-surgical MRI-images were used to build a 3D model of the skull and cerebellum for anatomical localization of cerebellar oculomotor vermis (lobules VI and VII) and lateral cerebellum (crus I/II; **Fig. 1B** (Sendhilnathan et al., 2020). Single-unit recordings were obtained using tungsten glass-coated electrodes (1-2 MΩ, Alpha Omega Engineering, Nazareth, Israel) through a 23-gauge guide tube, which was inserted only through the dura. A motorized microdriver (Alpha Omega Engineering, Nazareth, Israel) with a 1-mm spaced grid was used to introduce the electrode and map the recording sites with a maximum resolution of 0.25 mm.

### Data Analysis Software

All analyses were performed off-line using custom scripts written in MATLAB (Mathworks, Natick, MA, USA).

### Eye movement analysis

Eye position was sampled at 350 Hz with an infrared video eye tracker which tracked the pupil center of mass. Noise was reduced using a finite impulse response filter and a savitzky-golay filter (20 ms window). Eye speed and acceleration traces were created by differentiating the signal. Eye acceleration was further processed using a median filter. Saccade onset and offset were detected using an adaptive threshold based on 6 s.d. of the noise during fixation from the recording session of that day as described in Nyström and Holmqvist (2010) (Sendhilnathan et al., 2020). Trials with reaction times less than 80 ms were excluded, as these were thought to represent anticipatory actions.

### Electrophysiological analysis and statistics: Simple spikes (SSs)

To perform statistical analysis, neurons were required to have at least 5 recorded trials per direction as well as per pro- or antisaccadic condition. Only neurons with significant saccade-related SS activity and only correct trials were incorporated in the analyses. The latter choice was based on the fact that both NHPs Mo and Mi exhibited an expert level performance reaching >90% correct responses, rendering insufficient statistical power to analyze the incorrect trials. A neuron was considered saccade-related when SS firing during the baseline period (defined as the window 400 to 50 ms before trial onset in the intertrial interval), was significantly different from the saccade execution period activity (defined as the 150 ms time-window after saccade onset) for at least one of the 8 directions (Wilcoxon rank test; p < 0.05). We then computed the instantaneous SS firing rate of the neurons using a continuous spike density function (SDF) generated by convolving the spike train with a Gaussian function of σ = 50 ms width and averaging all the individual spike density functions (Sendhilnathan et al., 2020). We compared the number of spikes for all pro- and antisaccade trials of each PC using a Kolmogorov-Smirnov test, and subsequently determined whether the activity of a PC was significantly different between a pro- and antisaccade with a *p* value < 0.05. The characteristics of the saccade-related activity, as previously reported in PC recordings, was quite heterogeneous. On that account, we categorized the PC activity as *facilitation*, if the activity increased after instruction or saccade onset, or as *suppression*, if the activity decreased. For the instruction period, we compared the mean firing rate of each cell from a window 400 ms before instruction onset to a period of 300 ms at the end of the instruction period. For the saccade period we compared the mean firing rate of each cell from a window of 150 ms before saccade onset to a window from saccade onset to 150 ms after saccade onset (Wilcoxon signed rank, *p* value of < 0.05).

To determine the correlation between kinematic parameters and neuronal activity we determined Pearson’s correlation coefficient (*r*) for all neurons in the two tasks. Modulation ratios for both areas were obtained by computing the ratio between the response during pro- and antisaccades.

Changes in firing rate during the instruction period were computed by taking the differences between the mean firing rates during the baseline period separately for pro- and antisaccade trials. For the saccade period, we compared the mean activity 150 ms before the saccade onset to a period 150 ms after saccade onset.

We performed a demixed principal component analysis as described by Kobak et al. (2016) to decompose the population activity into individual components and extract the dependence of the PCs on the two stimulus conditions. The dPCA data analysis tool code is available at http://github.com/machenslab/dPCA (Kobak et al., 2016). Furthermore, all standard settings were used; data were aligned to instruction offset and binned in 100 ms bins. For determining cross-validated classification accuracies, 100 iterations were used.

### Electrophysiological analysis and statistics: Complex spikes (CSs)

Cells were included into the analysis for complex spikes if they had a CS firing rate of between 0.5 and 2 hz over the entire recording session and at least 5 trials for every of the 8 pro- and antisaccade directions. To determine if the CS modulated to one of the task epochs, we determined whether the CS rate significantly exceeded baseline levels during either the instruction or the saccade window (50 - 350 ms after onset of the instruction and −150 to +150 ms around the onset of the saccade). Baseline firing was calculated for a period of 500 ms of inter-trial activity, and the modulation was considered significant if firing rate exceeded ± 3 standard deviations from baseline.

Reciprocity of modulation between SS and CS rates was determined by finding the peak of the modulation of CS and SS responses during the same 300 ms windows (50 - 350 ms after instruction onset or −150 - 150 ms around the saccades). We first calculated the peak firing rate changes for SS and CS from baseline (ΔCS, ΔSS), and next determined the SS-CS interaction, i.e. reciprocity (ΔCS*ΔSS). Subsequently, we used a linear regression model to find out whether there was an association between reciprocity in the instruction period and reciprocity during the saccade period.

## Results

### Pro- and antisaccades and related PC activity in medial and lateral cerebellum

We trained two adult, male rhesus macaques (referred to as NHPs Mo and Mi) to perform a randomized, interleaved pro- and antisaccade task. In the prosaccade trials, the NHPs performed a (pro) saccade to a single visual target in one out of eight different locations separated 45° from each other. Instead, in antisaccade trials, the NHPs were trained to suppress the prepotent saccade to a visible target and perform a saccadic eye movement in the opposite direction to an unmarked, mirror position (**Fig. 1A**).

A trial started with the appearance of the central fixation point (instruction period), the color of which determined the trial condition. The colors red and green were the instruction cues for prosaccade and antisaccade movements, respectively (these colors are presented as magenta and blue in the figures to accommodate color-blind readers). Following the instruction period (300-500 ms), a target appeared randomly at one of the eight different locations, while NHPs maintained fixation on the central point. After 100 ms the central fixation point turned gray, serving as the “go-cue”. Animals had 500 ms to execute the pro- or antisaccade eye movement (saccade period) before the trial was aborted. This task design temporally separates the initial neural representation of the stimulus (300-500 ms long instruction period) from the subsequent movement execution (saccade period) (Amador et al., 1998).

Following the training period, we performed single-unit electrophysiological recordings. We recorded randomly from 90 PCs in medial and 72 PCs in lateral (Crus-I/II) cerebellum during 122 behavioral sessions (**Fig. 1B**). We removed trials with reaction times shorter than 100 ms following the “go-cue” to avoid contamination with data from “anticipatory” saccades; these reflect a visual, involuntary reflex towards a novel stimulus in the environment and do not represent correctly planned saccades (Goldring and Fischer, 1997; Coe and Munoz, 2017). We only considered correct trials for further analysis, as we did not have enough power to analyze error trials due to the high performance of both NHPs (94.1 ± 5.6% and 94.4 ± 5.3% for Mo and Mi, respectively). Eye movements during antisaccades exhibited longer reaction times, lower peak velocities, larger amplitudes, and longer durations than prosaccades (**Fig. 1C**).

A neuron was considered saccade-related if SS activity if activity during the intertrial interval (baseline period) was significantly different from the saccade execution period (i.e., from saccade onset to 150 ms after saccade onset) for at least one of the 8 directions (see **Methods**). Approximately half of the cells recorded in both areas fulfilled this criterion (n = 40 (44%) for the medial, and n = 38 (53%) for the lateral cerebellum). All subsequent analyses were performed only for these cells unless noted otherwise (n = 78 PCs). The average number of trials per cell in the medial cerebellum was 75 ± 49 s.d. and 71 ± 47 s.d. for pro- and antisaccades, respectively, while in the lateral cerebellum these numbers were 80 ± 52 s.d. and 76 ± 50 s.d., respectively.

To assess whether there was a correlation between saccade parameters and PC SS firing rate, we regressed the firing rates for each cell in the population (i.e. 90 PCs in medial and 72 PCs in lateral cerebellum) in the two recording areas to amplitude or peak speed and compared the corrected firing rate using the regression coefficient (ß) between pro- and antisaccade (corrected firing = firing rate/regression coefficient). We found no significant correlations between SS firing rate and saccade amplitude or saccade peak speed in neither the medial nor the lateral cerebellum (**Fig. 1D**). This allowed us to directly compare pro- and antisaccade trials between the two areas by pooling the mean firing rates of each cell in each condition. SS activity during saccade execution was significantly higher in the lateral cerebellum compared to that in the medial cerebellum for both pro- and antisaccade trials (mean and s.d., medial pro 66.7 ± 30 spks/s vs. lateral pro 69.3 ± 39 spks/s, *p* = 0.04; medial anti 68 ± 36 spks/s vs. lateral anti 71.8 ± 39 spks/s, *p* = 0.03; Wilcoxon rank sum test) (**Fig. 1E**).

### PCs in both medial and lateral cerebellum modulate their activity during the task but with different characteristics

The activity profile of the PCs followed roughly one of two patterns, they either increased or decreased their activity following saccade onset. Accordingly, we classified these cells as *facilitation* or *suppression* PCs, respectively (**Fig. 2**). The activity profile of the PCs was the same during pro- and antisaccades in both areas, meaning that if one PC increased its firing rate after saccade execution for prosaccades, it also increased it for antisaccades. This was true for all PCs except for two in the lateral cerebellum, which showed a mixed firing behavior; these two cells were included in their respective category for further analyses (e.g., as a *facilitation* cell in prosaccade trials and a *suppression* cell in antisaccade trials).

**Figure 2.**
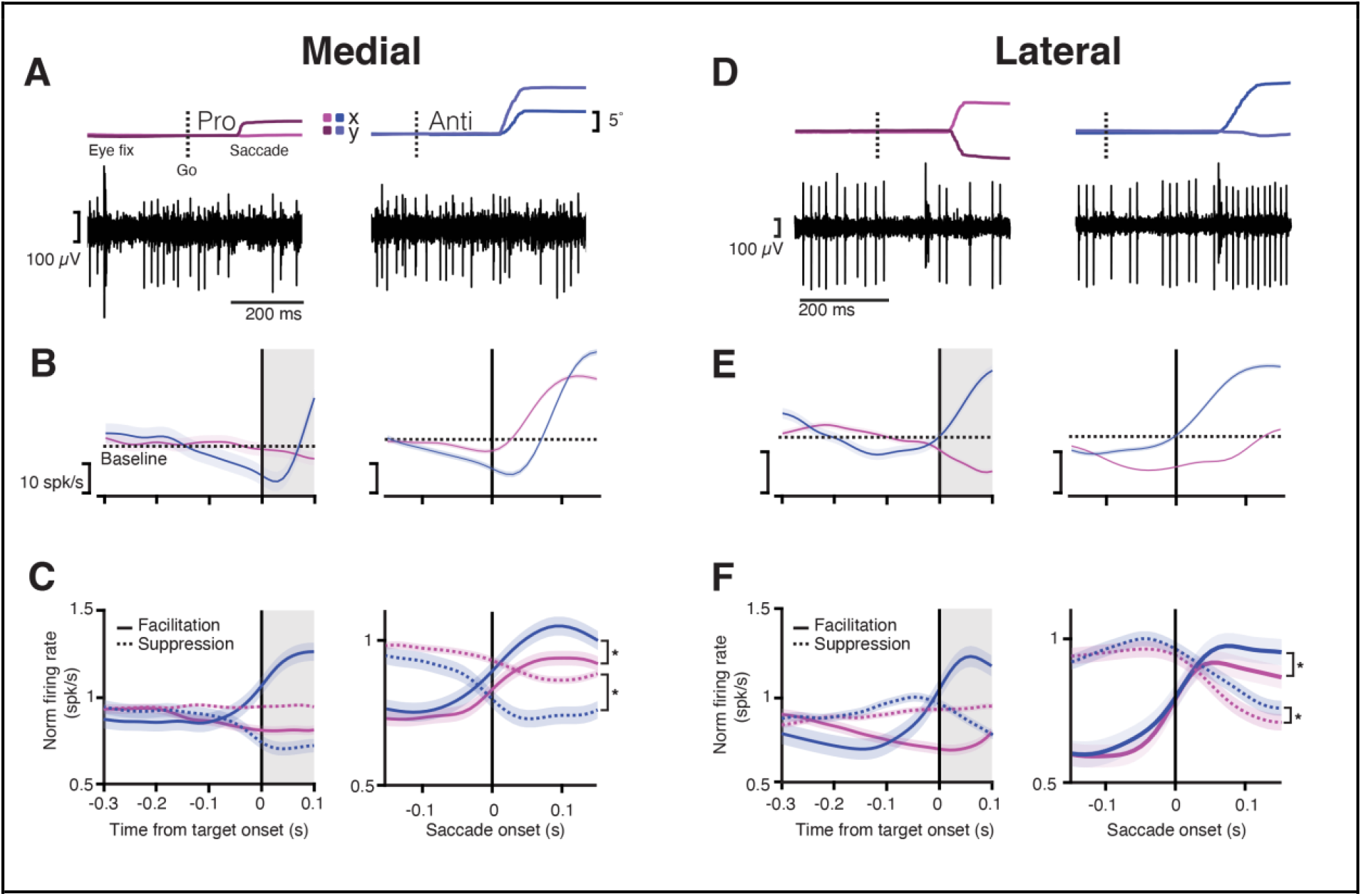
PCs in both medial and lateral cerebellum start modulating their activity during the instruction period. ***A top***, Eye trace for an example trial during saccade onset for a pro- (left) and antisaccade (right). Dashed line indicates the time of the “go” cue. Dark and light magenta and blue show the x and y position (horizontal and vertical components) of the eye trace for pro- and antisaccades respectively; ***bottom,*** traces of activity recorded from a PC in medial cerebellum for a single trial aligned to the eye trace indicated above. Note that the traces include many simple spikes (SSs; smaller spikes) and only a few complex spikes (CSs, bigger spikes). ***B***, ***left*** Average response time-course for one example neuron aligned to the end of the instruction period (target onset) for prosaccades (magenta) and antisaccades (blue) for medial cerebellum. Dashed line shows the baseline mean. Shaded regions denote s.e.m. Gray shaded areas show time after the instruction period when the target appears, but the animals must keep fixating at the central dot; ***right*** Average response time-course for one example neuron aligned to saccade onset. Dashed line shows the baseline mean. **C,** Time-course of normalized responses averaged for all neurons in medial cerebellum that either increased their firing (*facilitation* cells, solid line) or decreased it (*suppression* cells, dashed lines) aligned to target onset (left) or saccade onset (right). Statistical comparisons after saccade onset, facilitation pro vs anti: *p* = 1.61×10^-7^, suppression pro vs anti: *p* = 5.3×10^-23^. ***D***, Same as in (a) but for lateral cerebellum. ***E***, Same as in (b) but for lateral cerebellum. ***F,*** Same as in (c) for lateral cerebellum (facilitation pro vs anti: *p* = 1.08×10^-17^, suppression pro vs anti: *p* = 4.4×10^-9^). Prosaccades are denoted in magenta and antisaccades in blue for all panels.

We next assessed SS activity during the instruction period (for details, see **Methods**). Given the variable duration of the instruction period, which ranged between 300 and 500 ms, we analyzed the last 300 ms of this epoch, before target onset (**Fig. 1A**). Here, we also encountered diverse neural responses in both regions. We classified the activity of these neurons in the same way as for the saccade period, depending on whether they increased (*facilitation* PCs), or decreased (*suppression* PCs) their SS firing during the instruction period compared to the activity during the intertrial interval. The medial compared to the lateral cerebellum had overall less *facilitation* PCs (Medial pro 10%, medial anti ~11%, Lateral pro ~17%, Lateral anti ~18%) and *suppression* PCs (Medial pro ~11%, Medial anti ~7%, Lateral pro ~14%, Lateral anti ~15%) during the instruction of pro- and antisaccade trials (**Fig. 2B-F**). The remaining cells had no change in firing rates during the instruction period (medial: ~61%, lateral 36%).

To relate PC activity during the instruction period with the subsequent saccade period, we next looked at the timing of the SS activity of *facilitation* and *suppression* PCs between medial and lateral cerebellum pooling the pro- and antisaccade conditions (**Fig. 2**). We found that the latencies of the *suppression* PCs in the lateral cerebellum, measured at the trough after saccade onset, were significantly longer than those in the medial cerebellum (mean ± s.d. Lateral 133 ± 40 ms vs. Medial 89 ± 68 ms, *p* = 0.01; Wilcoxon rank sum, **Fig. 2C** and **F**). In contrast, the *facilitation* PCs showed similar timing of the peak activity in the medial and lateral cerebellum (mean ± s.d. Medial 105 ± 41 ms vs. Lateral pro 84 ± 48 ms; *p* = 0.4; Wilcoxon rank sum). There were no differences between prosaccades and antisaccades in this respect, neither for *suppression* PCs (mean ± s.d. Medial pro 87 ± 70 ms vs. Medial anti 93 ± 67 ms, *p* = 0.83; Lateral pro 139 ± 40 ms vs. Lateral anti 127 ± 42 s; *p* = 0.1; Wilcoxon Rank sum test), nor for *facilitation* PCs (mean ± s.d. Medial pro 105 ± 41 ms vs. Medial anti 104 ± 33 ms, *p* = 0.2; Lateral pro 84 ± 48 ms vs. Lateral anti 106 ± 43 ms, *p* = 0.9; Wilcoxon Rank sum test).

Together, these results suggest that PCs in the lateral cerebellum are more prominently involved in the instruction of planned saccades than those in the medial cerebellum, and that the course of SS responses of *suppression* PCs in the lateral cerebellum during the subsequent saccade period is delayed with respect to that in the medial cerebellum.

### PCs in lateral cerebellum show a stronger modulation during the instruction period

We observed that in some PCs, the SS modulation ramped throughout the interval between the instruction and the saccade epoch (**Fig. 2B-E** and **Fig. 3A**). We next tested if these saccade-related cells, also modulated their activity during the instruction period. For this reason, we quantified the ramping of SS activity by fitting a linear model to the firing rate during the last 300 ms of the instruction period, and we classified PCs as ramping cells when R^2^ was higher than 0.75. We found that ramping modulation was quite rare in the medial cerebellum and more prevalent in the lateral cerebellum (**Fig. 3B**); this held true for both prosaccade and antisaccade trials (Medial *n* (pro) = 3, *n* (anti) = 2; Lateral *n* (pro) = 11, *n* (anti) = 10; both comparisons *p* = 0.02; *χ*^2^ test). Overall, we found more PCs that significantly changed their firing from baseline during the instruction period in the lateral cerebellum than in the medial cerebellum (Medial vs Lateral, *p* = 0.03; Mann-Whitney test), suggesting a more prominent role for PC modulation in the lateral cerebellum during instruction.

**Figure 3.**
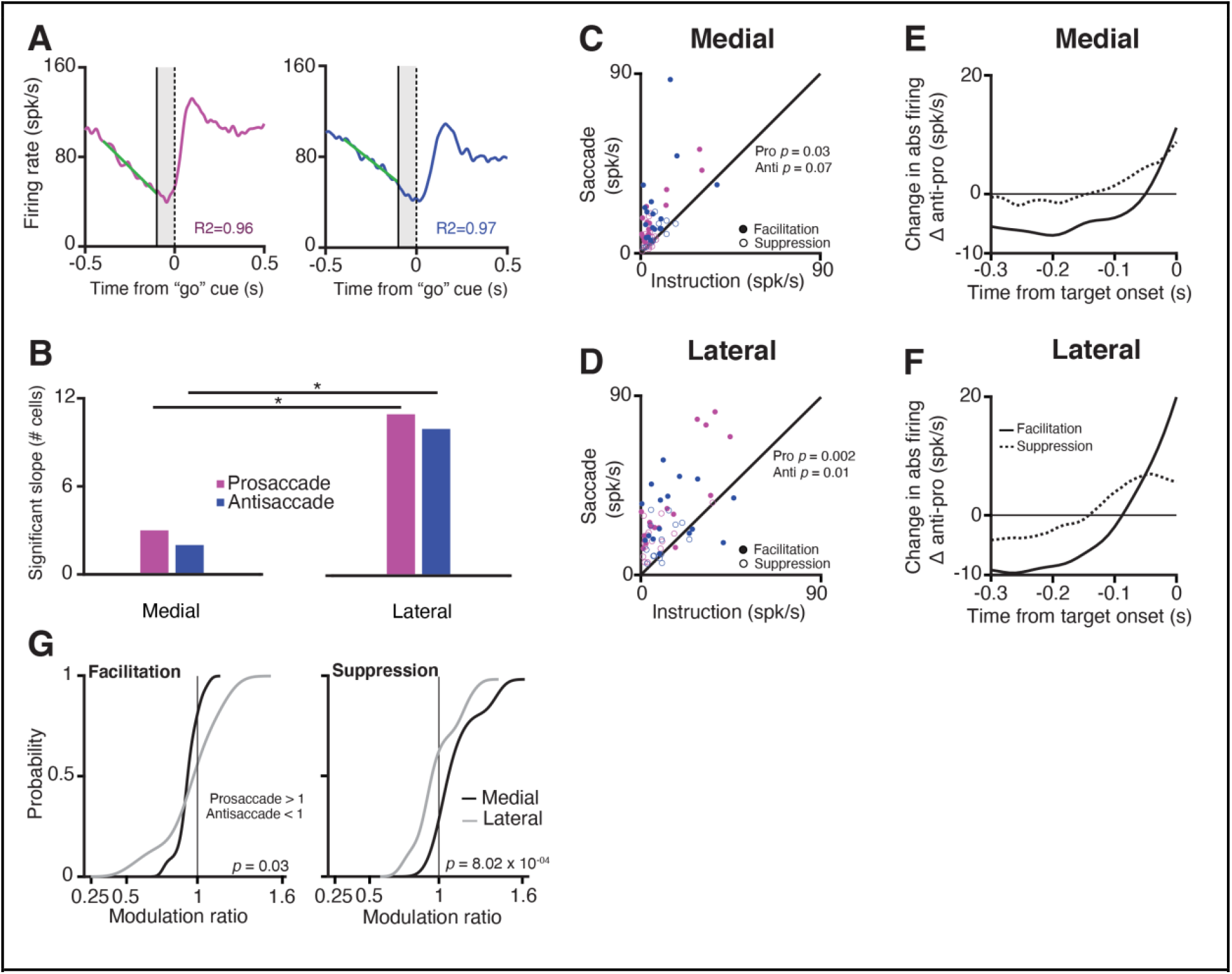
PCs in the lateral cerebellum exhibit stronger modulation than PCs in the medial cerebellum during the instruction period. ***A***, Explanation of slope calculations used for panel (b); ***left***, Time-course of responses during prosaccades (magenta) and antisaccades (blue) for an example neuron in the lateral cerebellum aligned to “go” cue (vertical dashed line at zero); *right*, activity of the same neuron, but now for an antisaccade trial also aligned to the “go” cue. Solid vertical line shows the end of the instruction period, shaded area is when the target appears but the NHPs must keep fixating. A linear regression model was fitted (orange diagonal solid and green dotted line, for pro- and antisaccades, respectively) to the firing rate 300 ms before the end of the instruction period to calculate the slopes (see **Methods**). ***B***, Number of cells with significant slope modulation in the medial and lateral cerebellum (χ2 test for proportions) during pro- and antisaccades (*p* = 0.02 for both comparisons). ***C***, Scatter plots of maximum simple spike firing rate change of PCs during the instruction period vs. that during the saccade period (in spks/s) for pro- and antisaccades in medial cerebellum. ***D***, Same as in (c) but for lateral cerebellum. ***E***, Difference in change in absolute firing rate between pro- and antisaccades (abs antisaccade SS activity-abs prosaccade SS activity) for *facilitation* and *suppression* neurons in medial cerebellum from target onset. ***F***, Same as in (e) but for lateral cerebellum. ***G,*** Cumulative distribution of the SS modulation ratio for medial and lateral cerebellum for *facilitation* and *suppressive* PCs. Insets show *p*-values for the comparison between medial and lateral cerebellum (two sample K-S test).

When we investigated the changes in the firing rate (modulation, **Methods**) for each PC during the instruction and saccade period within the medial and lateral cerebellum, we found that in the lateral cerebellum, the maximum change in the firing rate was significantly lower during the instruction period than during the saccade period, for both pro- and antisaccade trials (Lateral pro *p* = 0.002, anti *p* = 0.01; Wilcoxon rank sum test) (**Fig. 3C**). In the medial cerebellum only the prosaccade trails showed this trend (Medial pro *p* = 0.03, anti *p* = 0.07, Wilcoxon rank sum test) (**Fig. 3D**).

Further examination of the differences in responses during pro- and antisaccades showed that *facilitation* and *suppression* cells in the lateral cerebellum show significant encoding of SS modulation for trial conditions. We calculated a modulation ratio between the responses during the saccade period in the two types of trials, defined as the ratio between the response during pro- and antisaccades (**Fig. 3G**, **Methods**). Values of 1 indicate that both responses are the same, values higher than 1 indicate that prosaccade activity was higher, and below 1 that antisaccade activity was higher. Medial PCs had somewhat different responses when comparing pro- and antisaccade activity in contrast to lateral PCs (mean of *facilitation* PCs in medial cerebellum 0.91 ± 0.1 s.d. versus that in lateral cerebellum 0.88 ± 0.3, *p* = 0.03; mean of *suppression* PCs in medial cerebellum 1.16 ± 0.2 versus that in lateral cerebellum 1 ± 0.2, *p* = 8.02 x 10^-04^; one-tailed K-S test). To summarize, PCs in both medial and lateral cerebellum modulate their activity during instruction of pro- and antisaccades, with a relatively bigger role for the *facilitation* cells during the instruction of antisaccades. Yet, PCs in the lateral cerebellum differ from those in the medial cerebellum in that they have a relatively stronger modulation in both the instruction and saccade periods as well as for the pro-vs antisaccade conditions.

### Differences between facilitation and suppression cells that occur in both medial and lateral cerebellum

In addition to the observed differences across the recorded regions, we identified features that were shared across the medial and lateral cerebellum. First, we observed that *facilitation* and *suppression* PCs in medial and the lateral cerebellum operate at different baseline frequencies, which is in line with the upbound and downbound modules of PCs described for the cerebellar cortex of rodents (De Zeeuw, 2021). Since these modules have been hypothesized to operate at a relatively low and high baseline SS firing frequency, respectively, so as to leave ample room for increases and decreases during epochs of stimulation and modulation, we investigated whether the *suppression* and *facilitation* cells differed in this respect (see also (Sun et al., 2017; Herzfeld et al., 2018; Soetedjo et al., 2019). We found that *suppression* cells started at a relatively high baseline SS firing frequency in both the medial and lateral cerebellum (62.43 ± 3.76 spks/s and 69.6 ± 6.12 spks/s, respectively), and *facilitation* cells started at a significantly lower baseline SS firing frequency in both regions (56.36 ± 3.65 spks/s and 54.72 ± 3.76 spks/s, *p* = 0.02; one-tailed Wilcoxon rank-sum test comparison between all *facilitation* vs *suppression* cells pooled in both areas).

Next, we found that whereas the saccade period contained more *suppression* PCs, during the instruction period the contribution of the *facilitation* cells appeared most prominent, particularly during that of the antisaccade trials. When we subtracted the SS modulation during the prosaccade trails from the activity during the antisaccade trials (antisaccade SS activity - prosaccade SS activity), *facilitation* PCs in both medial and lateral cerebellum showed a larger change in firing rate just before the target onset during the instruction period (**Fig. 3E, F**). These changes were significantly less pronounced for *suppression* cells (*facilitation* PCs versus *suppression* PCs in medial cerebellum *p* = 4.4×10^-6^; *facilitation* PCs versus *suppression* PCs in lateral cerebellum *p* = 0.004; Wilcoxon Rank sum test). Thus, *facilitation* and *suppression* cells can be characterized by several features that are common for medial and lateral cerebellum.

### The lateral but not the medial cerebellum contains information about stimulus identity during the instruction period

To extract the features of the population activity related to pro- and antisaccades we applied a linear dimensionality reduction technique: demixed principal component analysis (dPCA, (Kobak et al., 2016). dPCA demixes the neural activity into activity related to the task components that capture the most variance in the data. The dPCA was used to investigate to what extent the neurons are tuned to two stimulus conditions indicated by the color of the fixation point (i.e. the pro- or antisaccades), or whether their activity is condition-independent. In the condition-independent group, dPCA components show task related activity modulations that cannot be assigned to one of the stimulus conditions. As explained above, due to the high behavioral performance of both NHPs, our dataset lacked the power to create another category with decision-related components in correct/incorrect trials. dPCA was applied to the entire population of recorded cells in the medial (n = 90) and the lateral cerebellum (n = 72). We included cells which had at least 5 trials in pro- and antisaccade conditions. The results were cross-validated to measure time-dependent classification accuracy and a shuffling procedure was applied to assess whether classification accuracy was significantly above chance level (see **Methods**).

In the medial cerebellum, the overall variance explained by the dPCA components was mostly captured by the first 3 components, which were independent of the stimulus condition, and together they explained 66.9 % of variance, (**Fig. 4A**). These components all have strong activity after the end of the instruction, and thus around the time the animal initiates the saccade (**Fig. 4B**). Components 4, 5 and 7 were selected as containing the strongest activity differences in relation to the stimulus conditions (**Fig. 4B**, bottom row and **Fig. 4C**). To determine significance, each component was used as a linear decoder to classify each of the stimulus conditions (see **Methods**). No components could reliably be used to classify the stimulus condition in the medial cerebellum.

**Figure 4.**
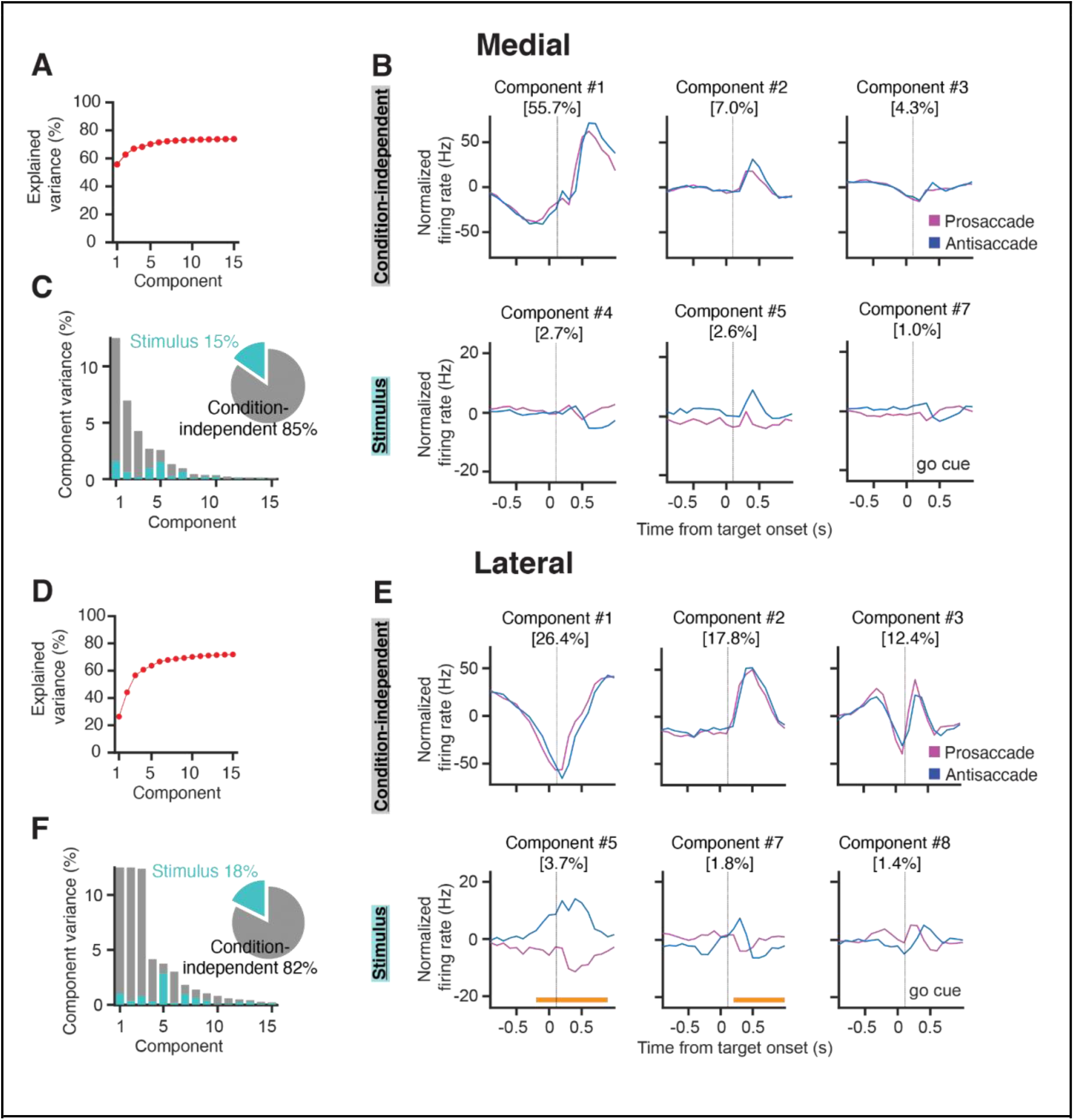
Demixed PCA indicates that PCs in the lateral cerebellum encode stimulus identity. ***A,*** Cumulative fraction of variance explained by dPCA in the medial cerebellum. Dotted line indicates an estimate of the fraction of variance in the data explained by the dPCA. ***B,*** Projections of the PSTH’s of all medial cerebellar PCs onto the most prominent decoding axes; ***top,*** first three condition-independent components; ***bottom,*** first three stimulus-related (pro/anti) components. Percentages denote variance captured by each component. Time from instruction offset. ***C,*** Variance of the individual demixed principal components for the medial cerebellum. Each bar shows the proportion of total variance. Gray bars depict condition-independent variance, cyan bars indicate stimulus-related (pro/anti) variance. Pie chart shows the total signal variance for stimulus-related and condition-independent components. ***D-F,*** Same as in (A-C) but for the lateral cerebellum, orange horizontal bars in ***E*** indicate time during which the stimulus can be decoded from the population firing rate.

Similar to the medial cerebellum, components 1, 2 and 3 of the lateral cerebellum captured most of the variance in the condition independent categories (56.4%, **Fig 4E**). These components also showed a strong modulation after instruction offset. In contrast to the medial cerebellum, two of these components modulated during most of the instruction period (**Fig. 4F).** Stimulus related components showed significant tuning for pro- and/or antisaccades in components 5 and 7. Notably, in component 5, stimulus condition classification was already possible almost 200 ms before the end of the instruction period. In the lateral cerebellum the variance in this component could almost be entirely attributed to the stimulus conditions (**Fig. 4F**). Indeed, in component 5, the time period in which the stimulus condition could be decoded from the firing rates was broad, bridging the instruction period to the saccade. This appears to be in line with the prominent ramping modulation described in **Figure 3**. These results show that PCs in the lateral, but not in the medial, cerebellum contain reliable information about the trial identity during the instruction period.

### Complex spike responses of PCs in the lateral cerebellum show reciprocal activity with simple spike responses

In a subset of recorded PCs, we were able to reliably isolate and analyze CS responses throughout all trials. We restricted the CS analysis only to cells with average CS firing rates higher than 0.5 Hz over the duration of the whole recording (Medial n = 11, Lateral n = 18). Similar to the SS analyses, we examined the CS responses separately in the medial and lateral cerebellum during the instruction and saccade periods (**Fig. 5**). The average CS activities during the instruction (1.10 Hz ± 0.47 s.d., n = 11) and saccade (1.11 Hz ± 0.50 s.d., n = 11) period in the medial cerebellum were not significantly different from those in the lateral cerebellum (1.15 Hz ± 0.33 s.d., n = 18 and 1.13 Hz ± 0.52 s.d., n = 18, respectively; *p* = 0.87 and *p* = 0.36 for comparison medial versus lateral during instruction and saccade period, respectively; Wilcoxon rank sum).

**Figure 5.**
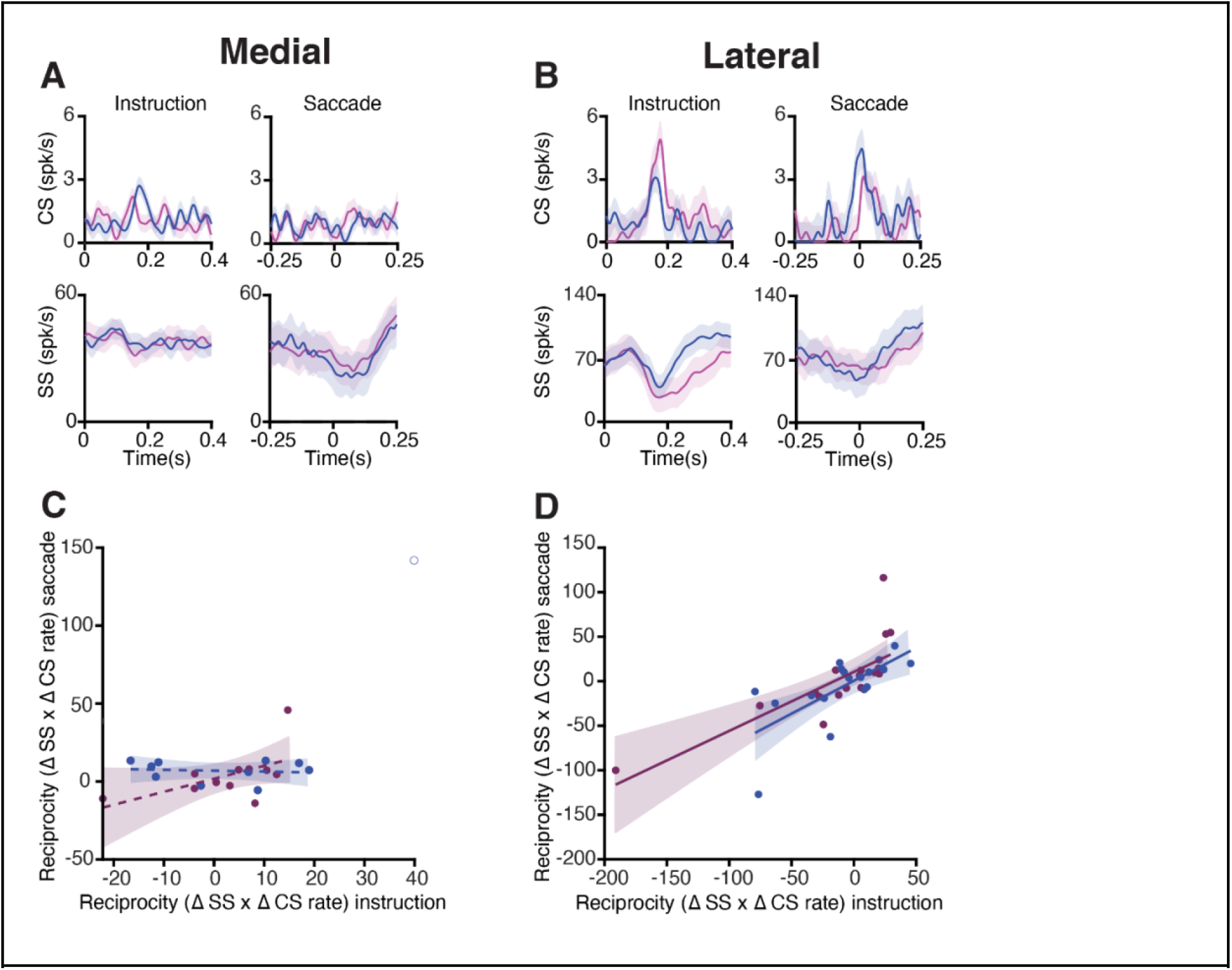
Complex spike and simple spike interaction during pro- and antisaccade trials. ***A***, Top, average CS rate of PCs in medial cerebellum during prosaccades (magenta) and antisaccades (blue) aligned to the onset of the instruction (left) and saccade (right) period. Bottom panels show the associated SS responses. Shaded areas represent s.e.m*. **B***, Same as *A*, but for neurons recorded in the lateral cerebellum. ***C***, Relation of reciprocity between CS and SS responses during instruction and saccade period. To quantify the reciprocity, we calculated the maximum change in CS firing rate in the windows of interest (instruction and saccade period), the associated maximum change in SS rate in a window of 300 ms around the peak of CS activity, and subsequently the product of the two (Δ CS rate x Δ SS rate). Note that if a positive modulation of CSs is associated with a negative modulation of SSs, the reciprocity measure presented here will give a negative value. If CSs and SSs show simultaneous positive modulations, and thus no reciprocity, this will result in a positive value. Every dot represents one cell. One outlier was removed from the medial antisaccade group (unfilled blue dot). The coefficients of determination (R^2^) are: medial pro 0.30, medial anti 0.01. **D**, same as C but of lateral neurons, coefficients of determination: lateral pro 0.59, and lateral anti 0.48. Shading indicates 95% confidence interval of the regression lines.

Next, we quantified the interaction between CS and SS responses during the instruction period (**Fig. 5A** and **B,** left panels). For every PC of both medial and lateral cerebellum we calculated the maximum change in CS firing rate from the baseline in the instruction window and the maximum change in SS firing rate from baseline in the 300 ms window around the time of the peak CS rate. This was also done for the saccade period for the pro- and antisaccade trials separately. Change in CS firing from the baseline was multiplied with change in SS firing from baseline to establish their interdependence, i.e., reciprocity (Badura et al., 2013). To determine whether reciprocity was dependent on both epochs, linear regression was applied on the reciprocity values in the instruction and saccade windows (**Fig 5C, D)**. Lateral cerebellar PCs had a positive association with reciprocity during the instruction and the saccade period, whereas medial cerebellum did not (coefficients of determination (R^2^): lateral pro 0.59, and lateral anti 0.48, medial pro 0.30, medial anti 0.01). Due to the relative lack of error trials, we unfortunately were not able to analyze error triggered CSs with sufficient power in a direction selective error context, as has been successfully done by other authors (Herzfeld et al. 2018).

## Discussion

The antisaccade task has been used to investigate flexible behavioral control, where a correct suppression of a prosaccade followed by a response to the opposite direction forms a feature of executive control (Mokler and Fischer, 1999; Mitchell et al., 2002). We found evidence for cerebellar involvement in volitional control of this behavior in both medial (i.e., OMV) and lateral (i.e., crus I/II) cerebellar cortex by recording the activity of PCs during pro- and antisaccades in NHPs. Our findings add to the growing body of evidence pointing to an important contribution of the cerebellum to cognitive processes in general and particularly voluntary behavior. We demonstrated that PCs in both areas modulate their activity depending on the trial type, and that SS responses of populations of PCs in the lateral cerebellum are able to discriminate the type of trial before the eye movement takes place. Even though the lateral cerebellum is not traditionally viewed as a saccade-control area, our findings support early reports where saccades are elicited following electrical stimulation to lateral regions of the hemispheres, especially in crus I/II (Ron and Robinson, 1973).

Antisaccades exhibit kinematic properties that differ from prosaccades, a result consistent with previous reports in human and non-human primates (Amador et al., 1998; Munoz and Everling, 2004). Even so, the preferential modulations of SS responses in the PCs were not correlated to any of those kinematic changes. This suggests that the observed activity was not merely an efference copy of the motor command but may also reflect the preparatory or planning phase of the voluntary movement in which the PC firing is determined by the cue. This is in line with recent findings in cerebellar atrophy patients who show impairments when making goal-oriented movements (Piu et al., 2019).

Various parameters of PC activity differed between medial and lateral cerebellum. SS activity of PCs in the lateral cerebellum (1) displayed more signal separation/selectivity for pro- and antisaccades during the instruction period, suggesting that the population contains information about the trial identity, (2) showed a greater firing rate during the execution of saccades. Likewise, their CS activity showed a higher level of reciprocity with respect to simple spike activity, during both the instruction and execution period. Together, these data indicate that different modules of different cerebellar regions can contribute to the same complex behavior, yet with different propensities.

### PC modulation during instruction period

Studies from cerebellar patients suggest that the cerebellum might not be involved in the initiation of saccades, but in the correct and accurate execution of them (Kornhuber, 1971; Thier et al., 2002). Our results indicate that PCs in the lateral cerebellum encode information about the trial identity. The dPCA analysis in Crus I/II PCs exhibited significantly different SS activity between both conditions, showing that these contain sufficient information to distinguish a pro- from an antisaccade. This is supported by a previous study showing that downstream neurons in the dentate nucleus modulate during preparation of antisaccades and that inactivation thereof promotes antisaccade errors (Kunimatsu et al., 2016). This instruction-related activity in antisaccade trials may be relayed to brainstem nuclei that are engaged in curtailing reflexive saccades (Everling et al., 1999). The persistent SS modulation of PCs in lateral cerebellum upon cue presentation suggests that it updates an internal model for well-timed activity during motor planning, which may be relayed to the prefrontal cortex (Gao et al., 2018).

Correct decoding of a visual stimulus is an essential component of the pro- and antisaccade task. Visual responses in the cerebellum have been reported in several areas including the floccular complex, crus I/II and the inferior semilunar lobe (VII/VIII) (Marple-Horvat and Stein, 1990). Dentate nucleus neurons have activity bridging the period from task instructions to motor execution (Kunimatsu et al., 2016), which could be derived from the same population of PC’s as we recorded in the lateral cerebellum. Our results on the relatively prominent CS - SS reciprocity in lateral cerebellum raise the possibility that CS responses play a similar role in decoding visual stimuli for the preparation of ensuing movements.

Contemporary models of antisaccade decision-making require evidence accumulation to reach a decision boundary in the frontal cortex (Munoz and Everling, 2004). PCs receive diverse sensory, motor and cognitive information through their parallel fiber inputs, which may facilitate efficient evidence accumulation. Classical models of cerebellar function indicate that climbing fiber mediated plasticity in the molecular layer determines which parallel fiber inputs are turned into SS modulation (Gao et al., 2012). The PCs that we measured in the lateral cerebellum carry both SS and CS signals during the instruction period of the antisaccade task. We hypothesize that through the pre-learned associations of parallel fiber and climbing fiber signals, relevant evidence from the stimulus (i.e., dot color in instruction) for the selection of the required action (i.e., pro- or antisaccade) can be rapidly relayed from the lateral cerebellum to the frontal saccade areas. In cerebellar patients, latencies for antisaccades are prolonged, indicating a less efficient decision-making process in absence of cerebellar input (Piu et al., 2019).

### PC modulation during saccade execution

Single PCs modulated their SS activity during the execution of pro- and antisaccades in both medial and lateral cerebellum, but only lateral PCs were able to successfully discriminate the type of trial. Despite the heterogeneity of the responses, neurons that facilitated or suppressed their response after saccade onset exhibited differential activity during pro- and antisaccade trials (**Fig. 3**). As a population, PC responses were different when executing a saccade towards a visible target than when using an internally generated goal. This is particularly interesting for two reasons: First, medial cerebellum has always been identified as a motor and timing controller for the different types of saccades, interacting with cortical and brainstem brain areas (McElligott and Keller, 1984; Noda and Fujikado, 1987; Thier and Möck, 2006; Herzfeld et al., 2015, 2018), while the role of the lateral cerebellum in saccade control has remained under debate. Indeed, lesions of lateral cerebellum can delay onset of saccades between 10 and 60 ms and result in variable hypo- and hypermetria (Ohki et al., 2009), but the contribution of Purkinje cell activity in the lateral cerebellum to proactive control of saccades and flexible behavioral in general has not been conclusively proven. Second, differential cerebellar activity during antisaccades implies that it contains neural signatures to modulate preparation and execution by encoding information about the current context of the upcoming action (saccade), which is necessary for the fast classification of the response as correct or erroneous. This finding is in line with the impact of focal cerebellar lesions in humans on the changes in potentials observed during performance monitoring in an antisaccade task (Peterburs et al., 2012).

### Cerebellar modules operate in parallel

The data on *suppression* PCs in the medial cerebellum showing a relatively early SS modulation during execution of both pro- and antisaccades as well as those on *facilitation* PCs in the lateral cerebellum showing a prominent modulation at the end of the instruction of antisaccades indicate that different modules of different cerebellar regions can simultaneously contribute to the same complex behavior, yet with different particularities. A similar conclusion was drawn from a recent study on delay eyeblink conditioning (Wang et al., 2020). Even though this form of conditioning is classically considered to be controlled solely by modules in lobule simplex of the lateral cerebellum, Wang and colleagues have shown that modules of the medial cerebellum are equally essential, yet also contributing in a slightly differential fashion, possibly regulating mainly muscle tone. Similar to the current pro- and antisaccade tasks, during eyeblink conditioning the medial and lateral cerebellum also both engage *facilitation* (*i.e., upbound*) and *suppression* (*i.e., downbound*) cells, and they also operate at relatively low and high baseline firing frequencies, respectively (ten Brinke et al., 2015; Wang et al., 2020). Our current finding that *facilitation* and *suppression* cells appear to play a more dominant role during the instruction and execution of the saccade task, respectively, further highlights the differential functional relevance of the upbound and downbound modules (De Zeeuw, 2021).

Previous reports of activity in the dentate nucleus, which is the main target of PCs in the lateral cerebellum, have shown mixed evidence regarding the direction of modulation during saccade-related behavior (Ashmore and Sommer, 2013; Kunimatsu et al., 2018). Kunimatsu and colleagues (2018) found only upward modulating units during self-initiated saccades, whereas Ashmore and Sommer (2013) revealed groups of both facilitating and suppressing cells during delayed saccades. Possibly, these differences reflect differences in the behavioral paradigms and/or downstream pathways involved (De Zeeuw, 2021). In our dataset, the modulation exhibited by PCs is bidirectional in both medial and lateral cerebellum with populations of both *facilitation* and *suppression* cells.

## Acknowledgements

We thank Kor Brandsma and Anneke Ditewig for their excellent biotechnical assistance; Beerend Winkelman for providing the MATLAB code for spike sorting; Chris van der Togt for providing assistance with stimulus presentation; Mario Negrello for providing the MATLAB code for saccade detection; Geert Springeling, Peter Thier, Peter W. Dicke, Ruud Smith for their help with anatomical scans and 3D model building; and Kaushik J. Lakshminarasimhan for helpful comments in the analysis. This research was supported by FP7-C7 European Commission, the Marie Curie Initial Training Network ITN-GA-2009-238214, ENW-Klein, ZonMw (VIDI/917.18.380,2018), Netherlands Organization for Scientific Research, European Research Council (Advanced Grant and Proof of Concept Grant), Medical NeuroDelta Programme, Topsector Life Sciences & Health (Innovative Neurotechnology for Society or INTENSE), and the Albinism Vriendenfonds Netherlands Institute for Neuroscience. This manuscript previosuly appeared online as a preprint: https://doi.org/10.1101/2021.03.26.437236.

## Author contributions

CIDZ, AB, MF, NF and EA designed the study and analysis. EA and PR performed surgeries on the animals. EA and NF executed the experiments and EA, NF, AB and PJH analyzed the data. EA, NF, AB and CIDZ wrote the first draft. All authors edited the manuscript.

## Data and code availability

The data that support the findings of this study and code for analysis of in vivo eye movement and Purkinje cell recordings are available from the corresponding author upon reasonable request.

